# Cytotoxicity mechanisms and composition of the glyphosate formulated herbicide RangerPro

**DOI:** 10.1101/2021.11.18.469091

**Authors:** Robin Mesnage, Scarlett Ferguson, Francesca Mazzacuva, Anna Caldwell, John Halket, Michael N Antoniou

## Abstract

Understanding the nature of co-formulants and toxic effects of major glyphosate-based herbicide (GBH) formulations is considered a research priority. Indeed, the toxicity of the co-formulants present in GBHs have been widely discussed and the European Union recently banned the co-formulant polyoxyethylene tallow amine (POEA). We provide a foundation for the development of new environmental epidemiological studies by reporting the presence of the most commonly used POEA, known as POE-15 tallow amine, in the widely used US GBH RangerPro. In order to understand if POE-15 tallow amine is present in RangerPro at a concentration at which it can exert toxic effects, we also tested the cytotoxicity of this GBH compared to glyphosate and POE-15 tallow amine in the human epithelial cell line Caco-2, a representative of the human intestinal epithelium, and the first to be exposed from the human diet to glyphosate herbicides. The lethal concentration 50 for each of these substances was 125 μg/ml, 17200 μg/ml, and 5.7 μg/ml, for RangerPro, glyphosate and POE-15, respectively. The Caco-2 cell cytotoxicity assay indicated that RangerPro is more cytotoxic than glyphosate, suggesting that its toxicity can be due to the presence of the POE-15 surfactant. RangerPro and POE-15 tallow amine but not glyphosate exerted cell necrotic effects, but did not induce oxidative stress. We show that RangerPro contains POE-15 tallow amine at a concentration at which it could exert toxic effects, which offers a starting point for conducting surveys of co-formulant exposure in human populations.

## Introduction

Glyphosate, an N-phosphonomethyl-derivative of glycine, is used as an active ingredient in herbicides, such as Roundup, to control weeds in agricultural fields and urban environments, but also to desiccate crops shortly before harvest. Glyphosate-based herbicides (GBHs) do not only contain glyphosate in water, but also other ingredients called co-formulants, which are listed a “inert” by manufacturers as they are deemed not to have target herbicidal action. The major co-formulants entering into the composition of Roundup herbicides are surfactants, which are used to allow glyphosate penetration through the waxy surface of plant leaves (Mesnage et al. 2019).

There are a number of routes through which humans can be exposed to the co-formulants present in GBHs. First, application of GBHs in agricultural, urban and domestic settings can lead to exposure via inhalation and dermal absorption. Second, foodstuffs, especially those derived from crops sprayed with GBHs just prior to harvest (e.g., cereals), could lead to dietary ingestion of co-formulants. Consequently, it is relevant and important to assess the toxicity of the complete formulated glyphosate herbicides and not just glyphosate alone to understand their health implications. However, a major obstacle to such investigations is that surfactants used in the manufacture of GBHs such as Roundup are generally considered as a trade secret or confidential business information. Although the full composition of a GBH has to be submitted to regulatory agencies, it does not always appear on the safety information sheets provided to applicators and consumers.

Understanding co-formulant composition of major Roundup formulations is considered as a priority by the European Union (EU) human biomonitoring programme consortium to monitor their effects in human populations (HMB4EU 2018). The HMB4EU consortium referred to POEA as a priority substance to evaluate exposure to glyphosate-based herbicide co-formulants (HMB4EU 2018). In 2016, the EU Commission recommended to the Member States a ban on the use of the co-formulant POE-tallow amine from GBHs. In a previous study, we developed and validated a mass spectrometry method to measure the urinary excretion of surfactants present in Roundup MON 52276, the European Union GBH representative formulation (Mesnage et al. 2021a). We validated the method and showed that it is highly accurate, precise and reproducible to estimate the oral absorption of MON 52276 surfactants in Sprague-Dawley rats exposed to this GBH via drinking water. In this report, we provide a foundation for the development of new environmental epidemiology studies by reporting the presence of the most commonly used POEA, known as POE-15 tallow amine, in the widely used US GBH formulation RangerPro.

Tissue culture cell assays have proven useful to provide a rapid evaluation of the relative acute toxicity of different GBH formulations (Mesnage et al. 2013). In a previous investigation, a comparison of 9 GBHs to glyphosate by combining mass-spectrometry and cell culture to identify the contribution of POEA to the toxicity of these products found that all the formulations were more cytotoxic that glyphosate alone (Mesnage et al. 2013). In this study we therefore also tested the cytotoxicity of RangerPro, which we compared to the toxicity of glyphosate and POE-15 tallow amine. We chose to undertake this investigation using the human epithelial cell line Caco-2 as a representative of the human intestinal epithelium, which is the first tissue exposed to GBHs through the diet.

## Materials and Methods

### Chemicals

Standard reagents were of analytical grade and obtained from Sigma Aldrich (Gillingham, Dorset, UK), unless otherwise stated. Glyphosate was obtained from Sigma Aldrich N (CAS: 1071-83-6, purity ≥ 96%, catalog no: 337757). The formulation RangerPro was sourced from the US market. POE-15 tallow amine was purchased from ChemService (distributed by Greyhound Chromatography and Allied Chemicals, Birkenhead, UK). Stock solutions of glyphosate, RangerPro and POE-15 were prepared in FBS-free medium. RangerPro was diluted accordingly in phenol red-free Dulbecco’s Modified Eagle Medium (DMEM). DMEM without phenol red, foetal bovine serum (FBS), 10 μg/ml penicillin/streptomycin and trypsin (0.05 and 0.5%), 2 mM glutamine, penicillin/streptomycin and DMSO were all obtained ThermoFisher Scientific (ThermoFisher Scientific, Loughborough, UK.

### Cell culture

The Caco2 cell line isolated from a human primary colonic tumour, was obtained from the American Type Culture Collection (ATCC) and used between passages 46 and 66, grown in phenol red-free DMEM and supplemented with 10% FBS, 2mM glutamine and 10g/ml penicillin/streptomycin. Cells were maintained in 75cm^2^ flasks (Corning, Tewksbury, USA) under standard culture conditions of 37°C and 5% CO_2_. Cells were grown until a maximum 70% confluency, washed with 5ml PBS and incubated in 5ml 0.05% trypsin-EDTA) for 5 minutes to detach and then split as needed or seeded into 96 well plates to initiate experiments.

### Cytotoxicity assays

Caco2 cells were seeded at 20,000 cells/well in 100μl of medium in clear 96 well plates. Following a 24 hr incubation, cells were treated with the pesticides diluted accordingly to the desired concentrations in maintenance medium. After a further 24 hr incubation an MTT assay was performed to assess cell proliferation and thus cytotoxicity according to the manufacturer’s instructions. Cells were incubated for 2 hr in MTT ([3-(4,5-dimethylthiazol-2-yl)-2,5-diphenyltetrazolium bromide] solution at 1 mg/ml in FBS-free DMEM. The resulting formazan precipitate was then dissolved by addition of 100 μL DMSO and quantified spectrophotometrically at 560nm using the GloMax Multi Microplate Multimode Reader (Promega, Madison, USA). Cell viability was expressed as a percentage relative to the negative control of untreated cell samples.

### ToxiLight cell necrosis assay

Caco-2 cells were seeded at 10,000 cells/well in 50 μl of medium in white-walled, clear-bottomed 96-well plates and incubated for 24 hr. Subsequently, cells were treated with the concentrations of pesticides that produced LC50, the cytotoxicity threshold (LC99) or with the LC99/5. Following incubation, the Lonza Toxilight kit was used according to the manufacturer’s instructions. Briefly, a 50μl aliquot of the AK reagent was added to each well and after 5 minutes the plates were read using the GloMax plate reader with excitation 485nM and emission 520nM settings. The background luminescence from wells with media only was subtracted, and luminescence compared relative to the negative control, untreated cell samples. Triton X100 was used as a positive control, necrosis-inducing agent.

### Oxidative Stress

Caco-2 cells were seeded at 1×10^4^ per well in 80 μl medium in 96 well white-walled plates, incubated for 24 hr and then treated with the pesticides at the desired concentrations. The positive control, ROS-inducing agent treatment was with 50μM menadione. Immediately after treatment, 20 μl H_2_O_2_ substrate, was added to each well. At 6 hr post treatment, 100 μl ROS-Glo detection reagent (Promega, Southampton, UK) was added per well, as per manufacturer’s instructions. The plates were then incubated at room temperature in the dark for 20 minutes and the luminescence read using the Glo-max plate reader at excitation 485nM and emission 520nM settings.

### Mass spectrometry analysis

Formulation samples were diluted 1:100 with acetonitrile-water (1:1) containing 0.1% formic prior to their injection into the LC-MS/MS system. The UHPLC-MS/MS instrument consisted of a Thermo Scientific Accela™ UHPLC coupled to a Thermo Scientific TSQ Quantum Access™ mass analyser with a heated electrospray ionization source (HESI-II). The mass spectrometer was operated in positive mode and data acquired with Xcalibur software. The injection volume was 5 μL and the chromatographic separation was achieved using a Thermo Scientific Accucore C18 column (2.6 μm, 100 x 2.1 mm) maintained at 40°C. A binary gradient profile was developed using water with 0.1% formic acid (A) and acetonitrile with 0.1% formic acid (B) at a flow rate of 200 μL/min. HPLC grade acetonitrile and HPLC grade water were from Fisher Scientific, and LC-MS grade formic acid was from Merck (Merck Life Science UK Limited, Gillingham, UK). Separations were conducted under the following chromatographic conditions: 95% solvent A for 0.2 mins, decreased to 5% over 15 mins, maintained for 5 mins at 5% before being increased to 95% over 0.1 mins. Column equilibration time was 4.2 mins, with a total runtime of 25 mins. Mass spectrometric conditions were as follows: spray voltage 3.5 kV, capillary temperature 350°C, vaporizer temperature 300°C, sheath gas 50 au, auxiliary gas 5 au, full scan mode 150-1200 m/z.

### Statistical analysis

The dose response on Caco-2 cells was used to determine cytotoxicity thresholds using nonlinear dose-relationships. We determined the lethal concentration 50 (LC50), the concentration at which 50% of the cells are viable. Statistical analysis was performed by oneway ANOVA in GraphPad prism 5.

## Results and discussion

We determined the surfactant composition of RangerPro, a widely used herbicide in the US, using a mass spectroscopy approach. The comparison of RangerPro to the POE-15 tallow amine standard revealed that both products contain a complex mixture of surfactants (Figure 1A). Peaks for polyoxyethylene tallow amines with several different polyoxyethylene chain lengths were observed (Figure 1B). This is because the successive addition of ethylene oxide molecules to form POE does not occur at the same speed for each POE in the melange. When the chemical reaction is stopped, the POE chains produced have different lengths. Comparison of the m/z profiles of the two products confirmed that POE-15 tallow amine is a component of RangerPro.

**Figure 1.**
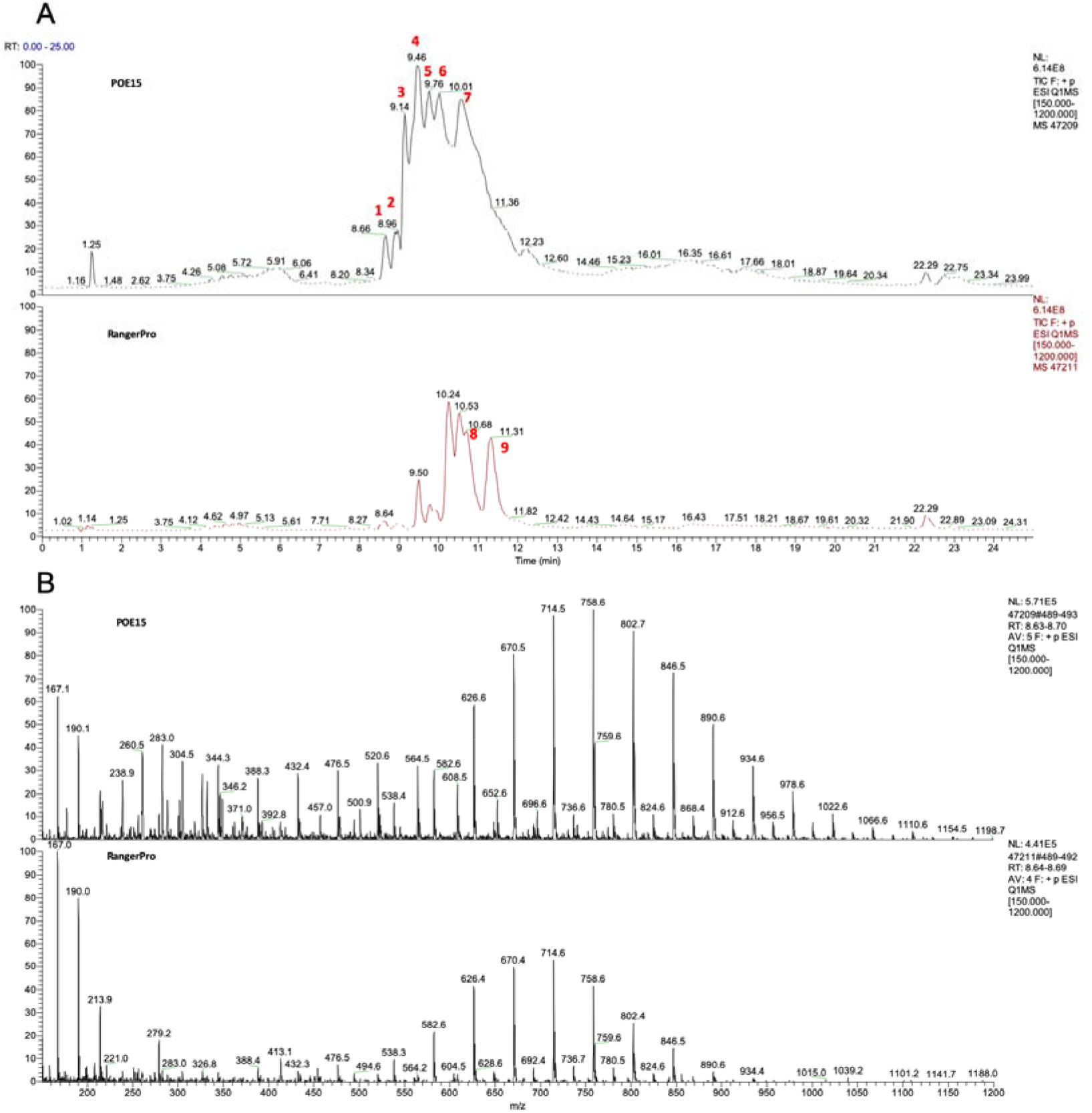
Representative spectrum of RangerPro obtained by liquid chromatography-mass spectrometry reveals the presence of POE-15 tallow amine. **(A)** Chromatograms of RangerPro and POE-15 show the presence of a mixture of co-formulants in the herbicide, which elute at different retention times from the mass spectrogram column. **(B)** The mass spectra (bottom panel) are obtained by extracting the TIC of peak 1, retention time: 8.7 min (TIC between 8.63-8.70 min) show that peaks are separated by 44 m/z increments, which is similar to what is seen with POE15 suggesting that each peak corresponds to a different molecule belonging to a homologous series of surfactants with a different number of ethoxylation units. Mass spectra from the other peaks 2-9 are available as supplementary data 1.

Human populations are exposed to surfactants from multiple sources, including pesticides but also washing products or cosmetics. It might thus not be fully clear whether the presence of surfactants in human bodily fluids originate from exposure to a pesticide. There is thus a need to conduct biomonitoring studies and assess for their presence in population groups, which are spraying pesticides containing surfactants, and correlate this exposure to glyphosate levels. Our demonstration that RangerPro contains POE-15 tallow amine, offers a starting point for conducting such surveys of co-formulant exposure.

We next investigated the cytotoxic potential of RangerPro compared to glyphosate alone to reveal the greater potential health risks from exposure co-formulants present in the herbicide formulation. A Caco-2 cell cytotoxicity assay was performed, which gave LC50 values of 125 μg/ml, 17,200 μg/ml, and 5.7 μg/ml, for RangerPro, glyphosate and POE-15, respectively (Figure 2). Thus, based on this assay RangerPro was ~22 times and POE-15 >3000 times more cytotoxic than glyphosate. This demonstrates enhanced toxicity from the presence of the POE-15 surfactant and possibly other related surfactants in the RangerPro formulation.

**Figure 2.**
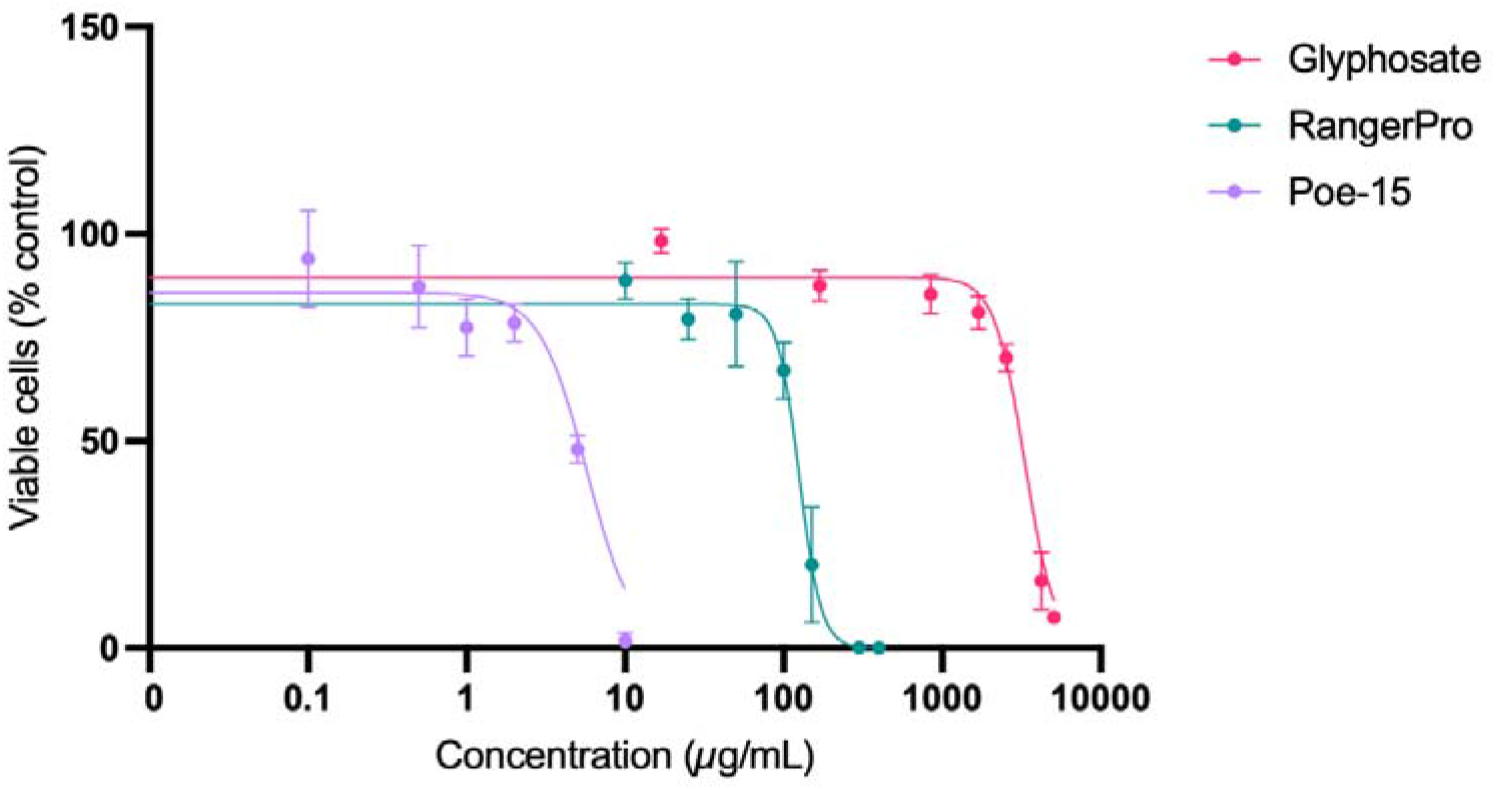
RangerPro is markedly more cytotoxic than glyphosate alone in Caco-2 cells. Caco-2 cells were treated with the test substances for 24 hr, and viability determined by an MTT assay. The concentration of RangerPro is expressed as glyphosate equivalent concentrations. Cell viability was expressed as a percentage relative to the negative control, untreated cell samples. The assay was performed in triplicate and data is expressed as mean ± SD (standard deviation) of 3 independent replicates.

A large number of studies have shown that POEA surfactants contribute to the toxicity of glyphosate herbicides. Studies showing that POE-15 tallow amine is more toxic than glyphosate stretch back to 1979 (Folmar et al. 1979). The formulation MON 2139 containing POE-15 tallow amine was 10 to 40 times more toxic than glyphosate in different fish species (Folmar et al. 1979; Wan et al. 1989). Further studies showed that the lethal concentration at which 50% of rainbow trout (*Oncorhynchus mykiss*) were killed was 86 mg/L for glyphosate, and 8.2 mg/L for MON 2139. The herbicide MON 2139 was also 10 to 50 times more toxic than glyphosate in four North American amphibian species (Howe et al. 2004; Mann and Bidwell 1999) or Microtox bacterium, microalgae, protozoa and crustaceans (Tsui and Chu 2003). In a previous investigation using three mammalian tissue culture cell lines (HepG2, HEK293, JEG3), we showed that 9 GBHs were up to 100 times more cytotoxic and POE-15 was 10,000 times more toxic than glyphosate alone (Mesnage et al., 2013). Thus, our results showing that RangerPro is 22 times and POE-15 >3000 times more cytotoxic than glyphosate alone in Caco-2 cells (Figure 2) is in accord with these earlier observations.

The mechanism of action by which POEA caused cytotoxicity in previous studies was by disrupting the structure of cell membranes causing necrosis (Mesnage et al. 2013). Therefore, we next undertook an analysis to see if RangerPro also caused cellular necrosis (Figure 3). We tested the LC50 (cytotoxic), LC99 and LC99/5 (non-cytotoxic) threshold concentrations. RangerPro and POE-15 treatment resulted in a statistically significant increase in adenylate kinase release indicative of induced necrosis at all concentrations tested (Figure 3). Glyphosate did not cause necrotic effects, confirming that the cytotoxic capability of RangerPro can be attributed to the membrane disrupting potential of the surfactants included as co-formulants.

**Figure 3.**
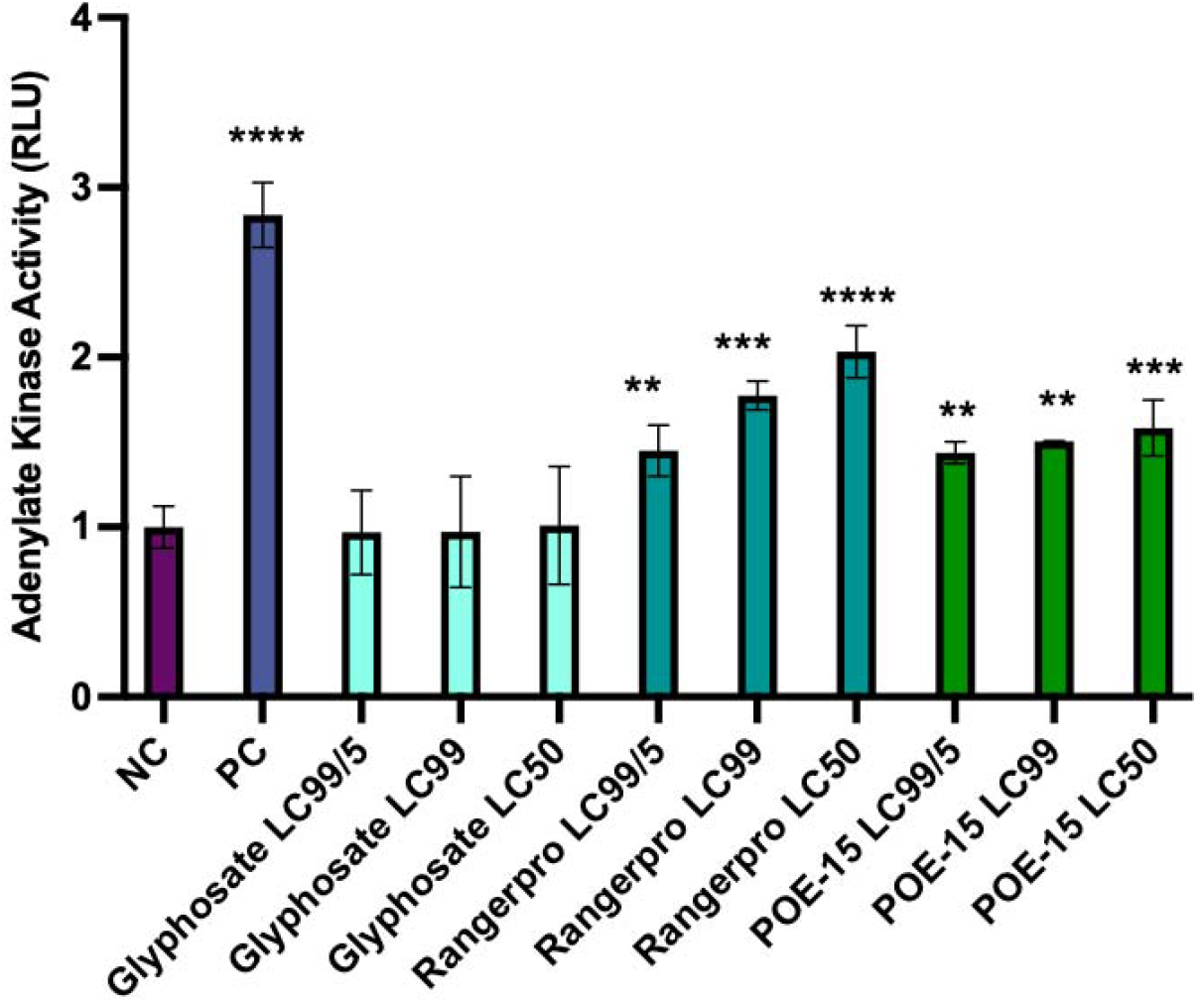
Cytotoxicity of Caco-2 cells by RangerPro and POE-15 is via a membrane disruption and necrosis mechanism. Caco-2 cells were treated with 3 concentrations of each test substance, which corresponded to the lethal concentration (LC) thresholds determined by the MTT cytotoxicity assay. The positive control (PC) was a concentration of Triton-X100 surfactant known to cause cell necrosis. The assay was performed in triplicate and data is expressed as mean ± SD (standard deviation) of at least 3 independent replicates. Adenylate kinase activity shown is represented as fold change in relative light units (RLU) relative to the negative control (NC) untreated cells culture samples. Statistical analysis was performed by one-way ANOVA in graphpad prism 5 (** p<0.01; *** p<0.001; **** p<0.0001).

Since glyphosate has frequently been described as a disruptor of redox status in mammalian cells (Mesnage et al. 2015), we tested to see if either glyphosate or RangerPro could increase production of hydrogen peroxide, an indicator of oxidative stress in Caco-2 cells (Figure 4). The positive control substance menadione provoked a significant (6-fold) increase in hydrogen peroxide production in comparison to the negative control, untreated cell samples (Figure 4, NC and PC values). Although some of the test compounds caused an increase in oxidative stress, effects were limited. Interestingly, oxidative stress seemed to be reduced by the test compounds. This is not surprising because oxidative stress has known hormetic properties. This phenomenon by which mild-induced stress can give rise to a positive physiological counter-response inducing maintenance and repair systems, has already described by our group after an exposure to a low-dose pesticide mixture (Mesnage et al. 2021b), and for other pesticide exposures on both target (Tang et al. 2019) and non-target (Calabrese and Baldwin 2003) species. This outcome can thus be interpreted as an effect of glyphosate on redox systems, which is compensated by cellular metabolism. It should be noted, however, the generation of oxidative stress can be time-dependent and it is not clear if our experimental design captures this glyphosate property of toxic.

**Figure 4.**
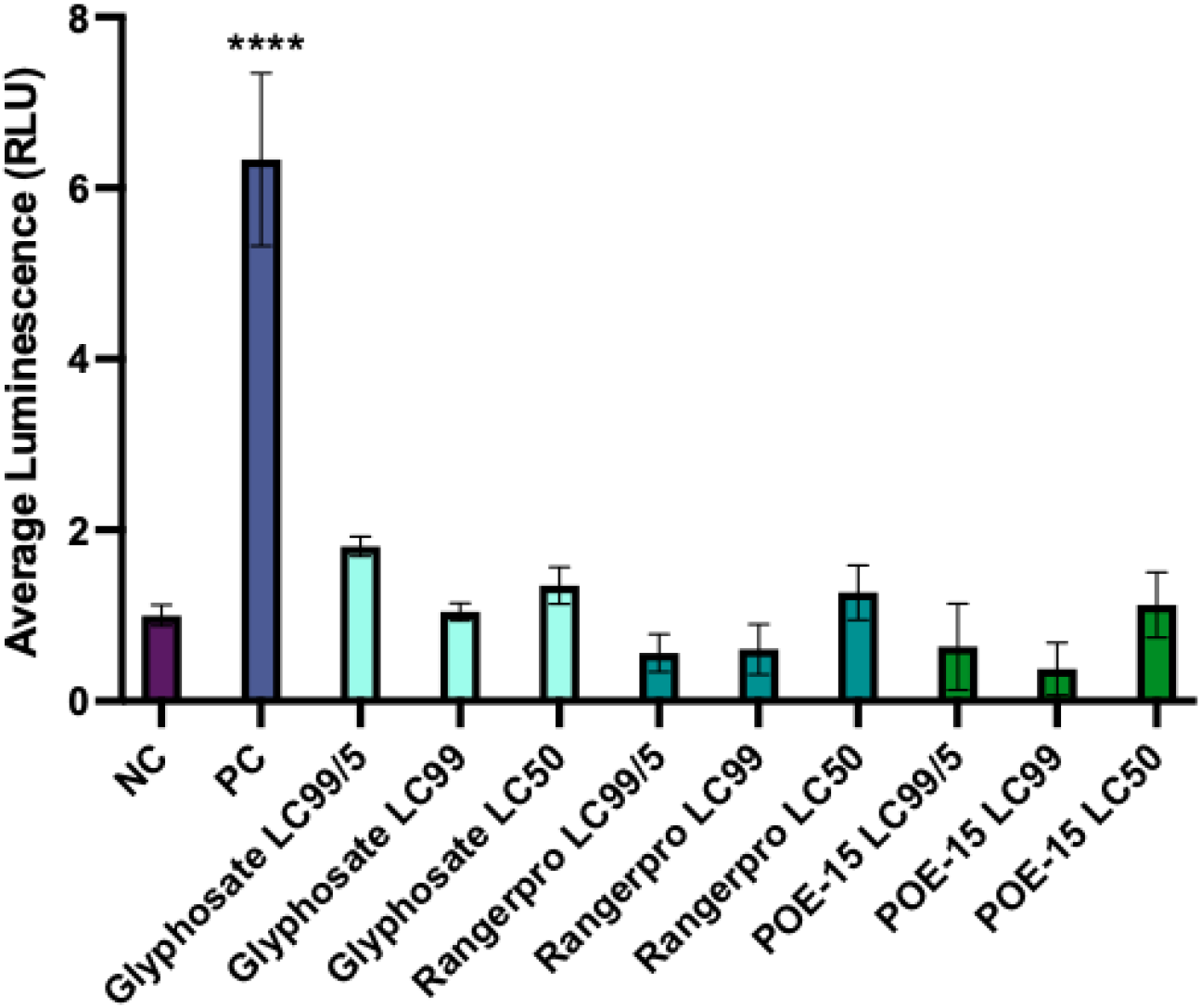
Oxidative stress measured by hydrogen peroxide production. Caco-2 cells were treated for 6 hr at respective LC50, LC99 and LC99/5 thresholds with glyphosate, RangerPro and POE-15. Treatment with 50 M menadione acted as positive control (PC). Production of hydrogen peroxide was the oxidative stress marker detected by the luciferase reporter system. The assay was performed in triplicate and data is expressed as mean ± SD (standard deviation) of 3 independent replicates.

Our study was performed on the human epithelial cell line Caco-2, chosen as a representative of the human intestinal epithelium. In our previous studies, we found that glyphosate inhibited the shikimate pathway in the gut microbiome, causing an accumulation of compounds upstream of the EPSPS enzyme (e.g. shikimic acid) (Mesnage et al. 2021c). Future studies would also need to take into account that the effects of glyphosate are multifactorial, and that glyphosate formulations might not only alter bacterial community composition but also damage the gut epithelium compromising its integrity.

## Acknowledgements

This work was funded by the Sustainable Food Alliance (USA) whose support is gratefully acknowledged.

