## Supplemental Data 1 for "Cytotoxicity mechanisms and composition of the glyphosate formulated herbicide RangerPro"

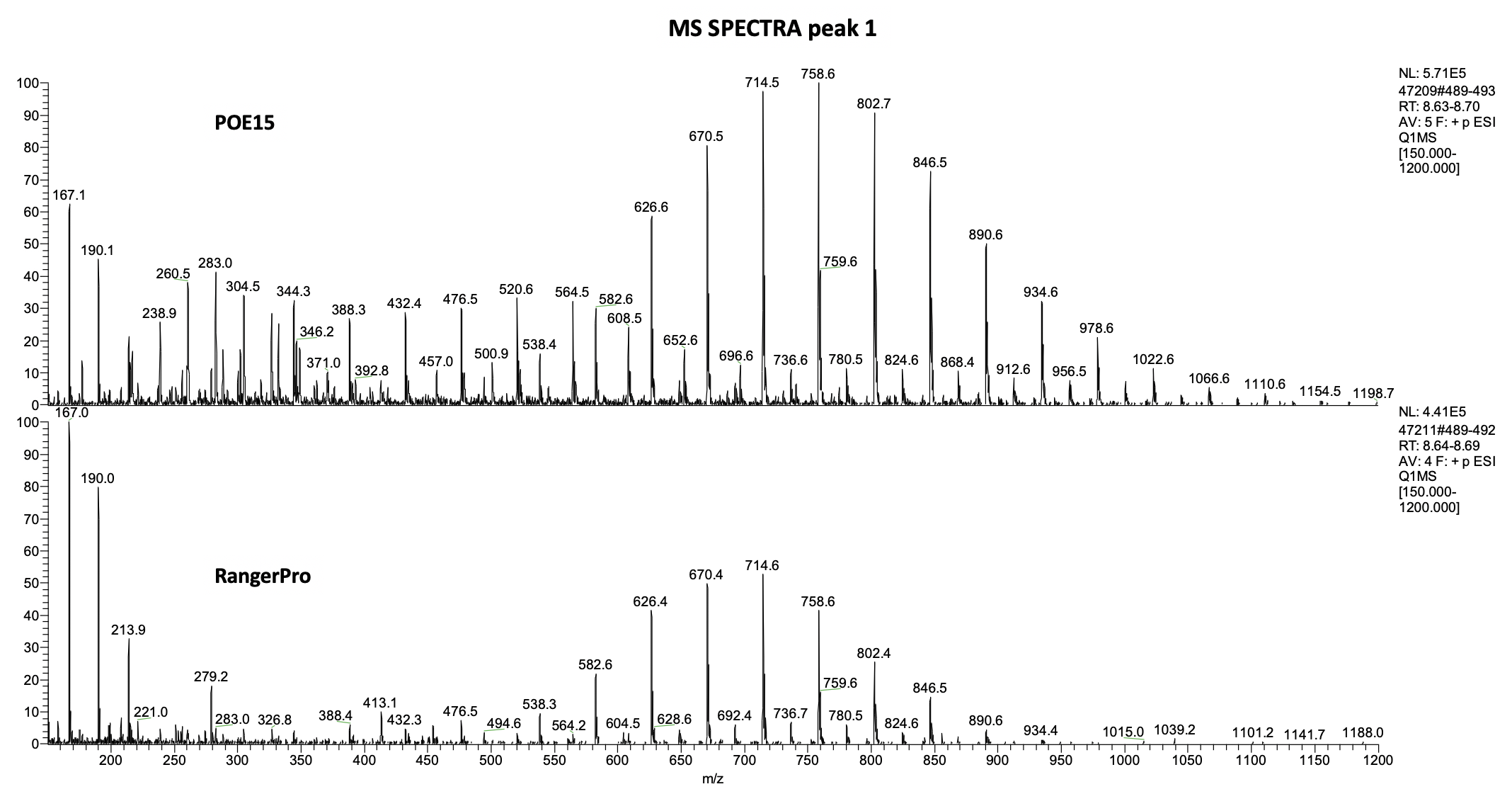

*Representative mass spectra obtained by extracting the TIC of peak 1 from the chromatogram of POE-15 and RangerPro which elute at different retention times (See figure 1).*

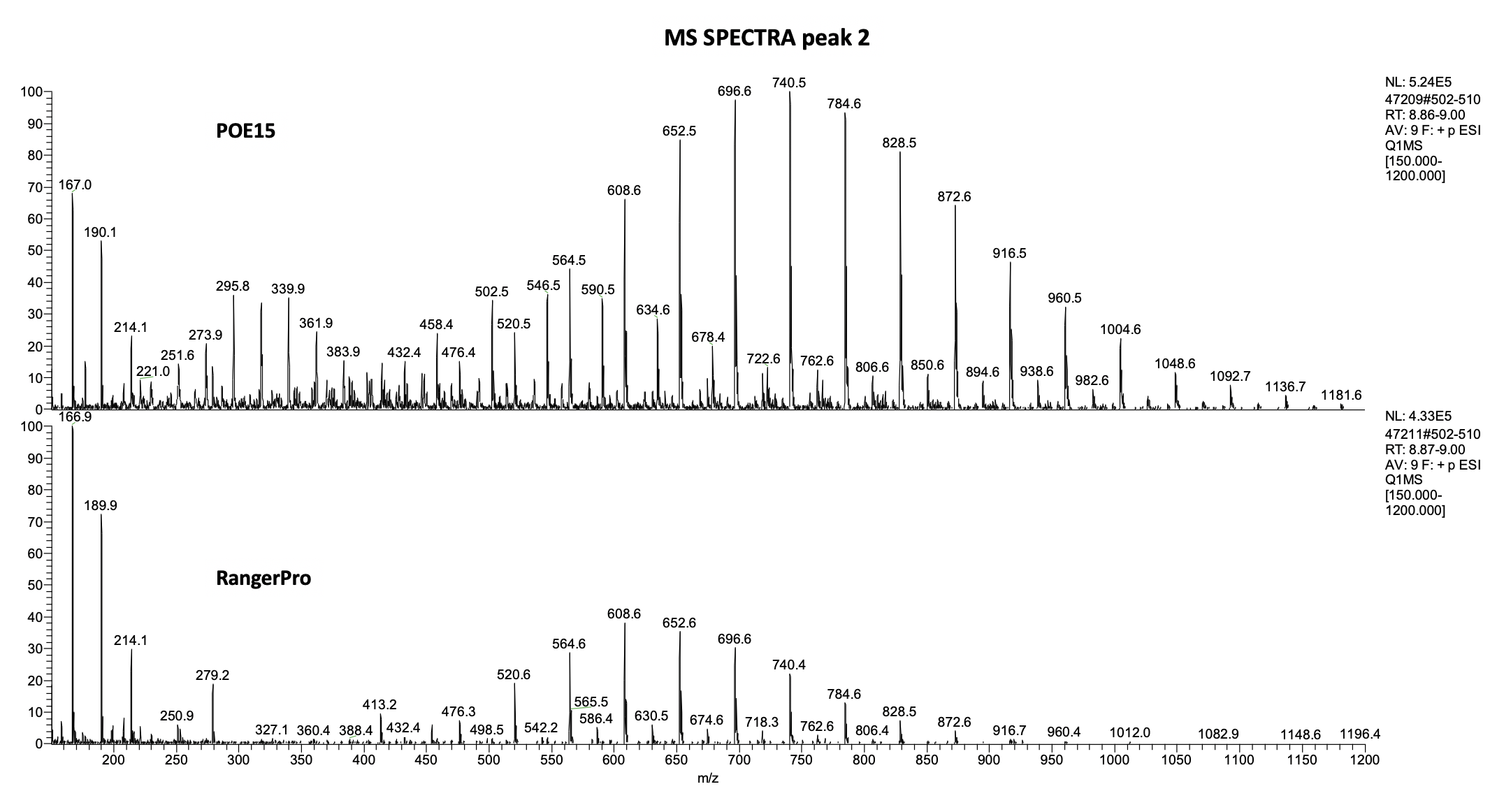

*Representative mass spectra obtained by extracting the TIC of peak 2 from the chromatogram of POE-15 and RangerPro which elute at different retention times (See figure 1).*

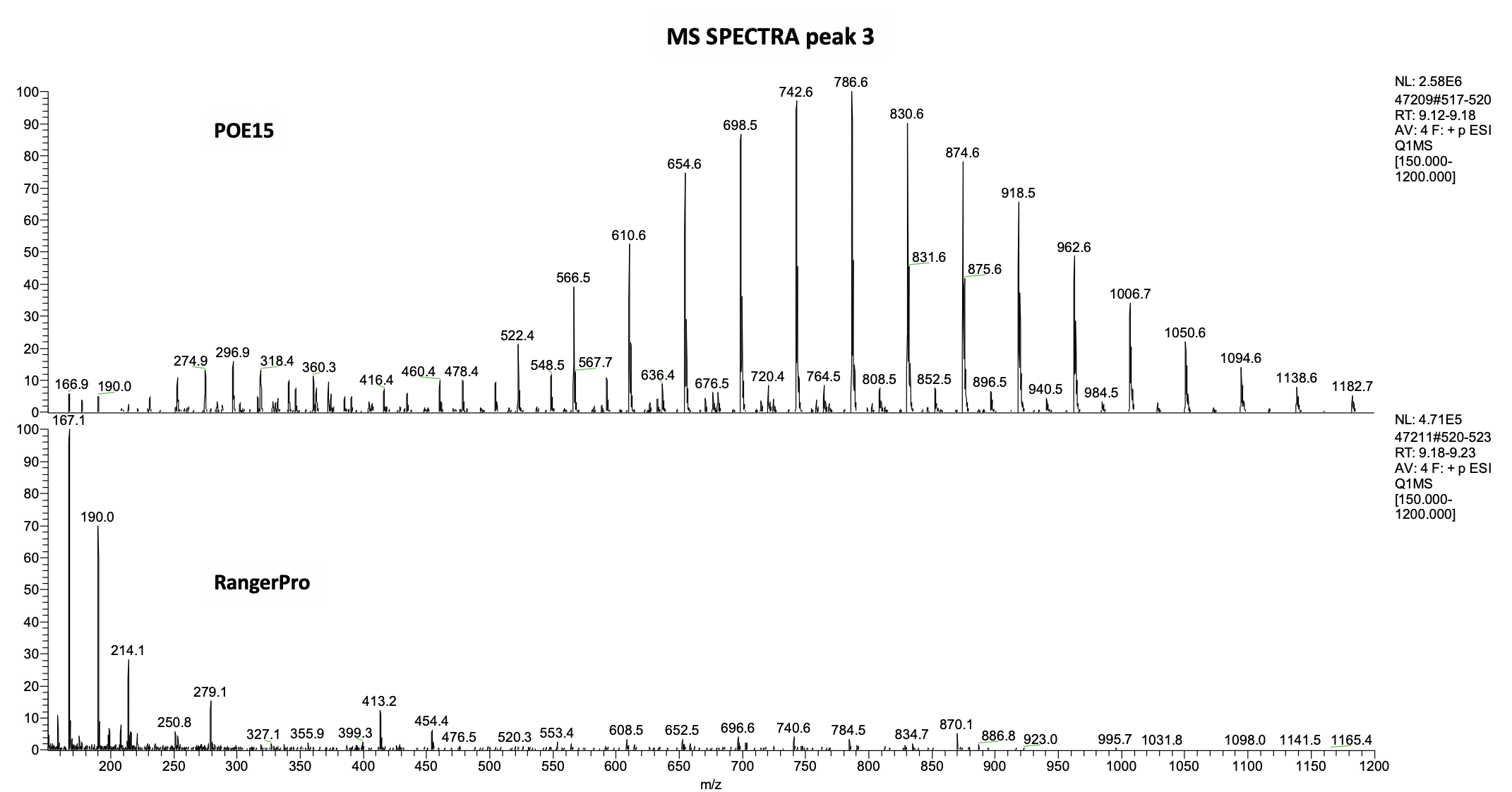

*Representative mass spectra obtained by extracting the TIC of peak 3 from the chromatogram of POE-15 and RangerPro which elute at different retention times (See figure 1).*

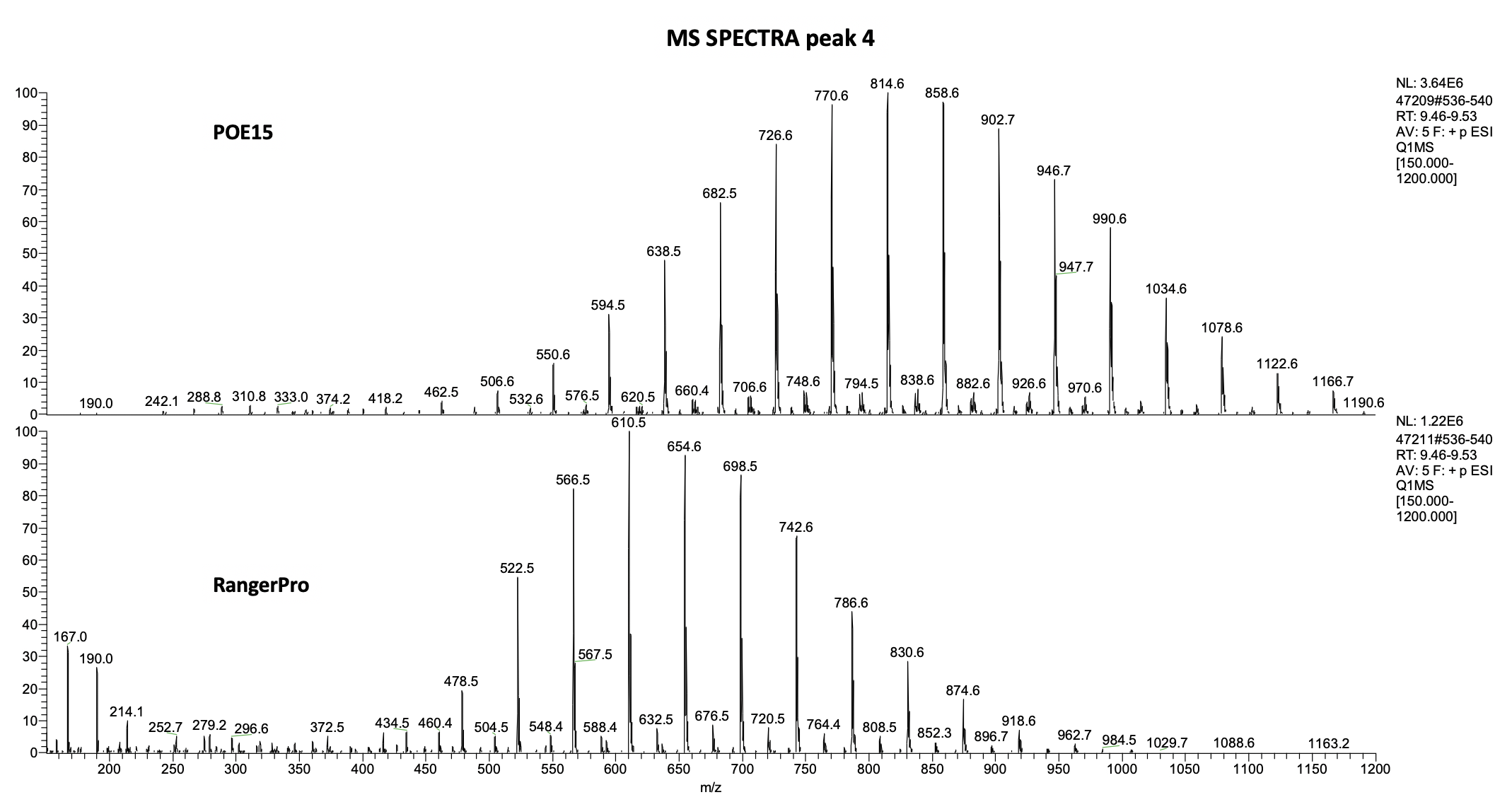

*Representative mass spectra obtained by extracting the TIC of peak 4 from the chromatogram of POE-15 and RangerPro which elute at different retention times (See figure 1).*

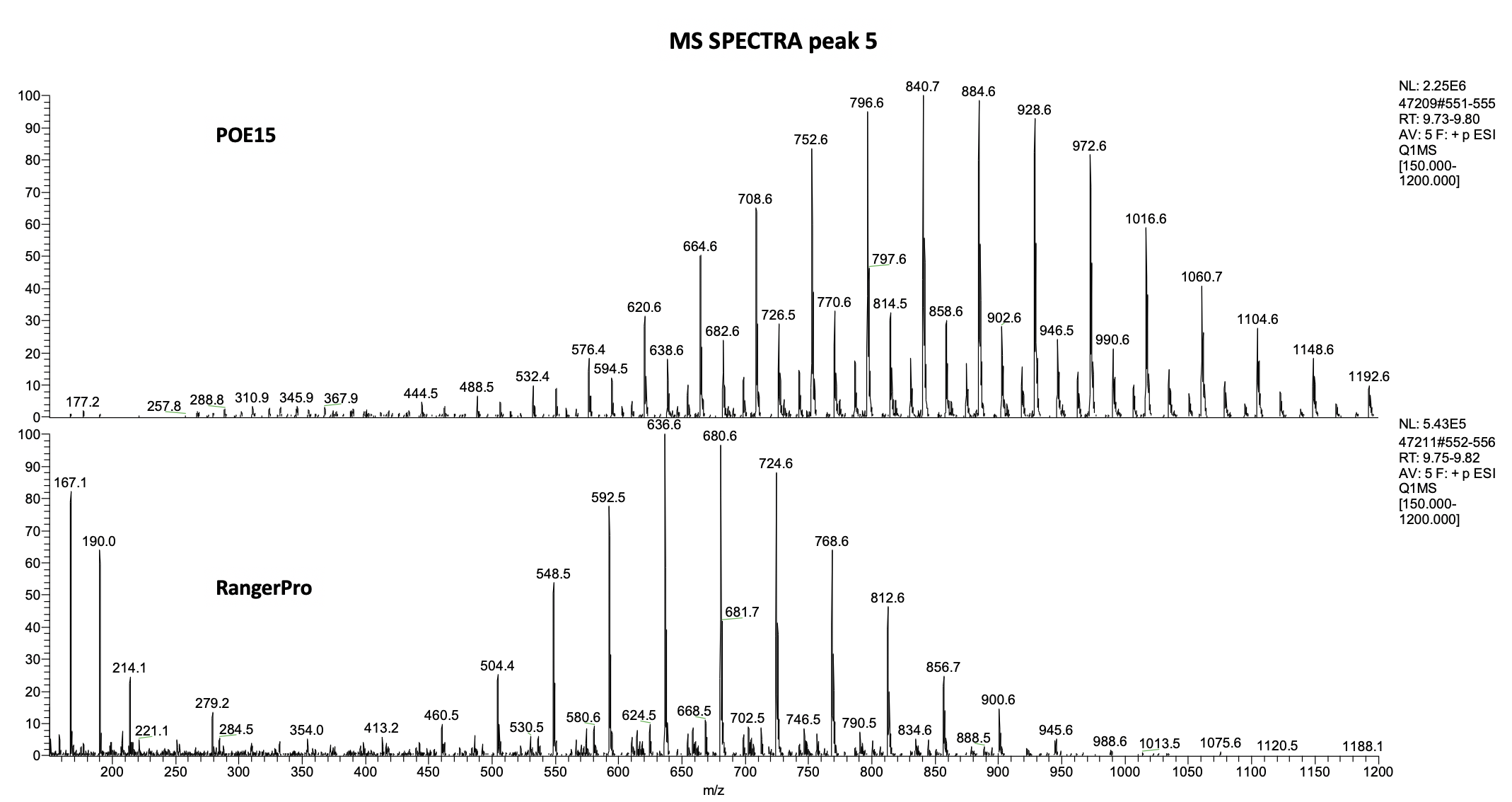

*Representative mass spectra obtained by extracting the TIC of peak 5 from the chromatogram of POE-15 and RangerPro which elute at different retention times (See figure 1).*

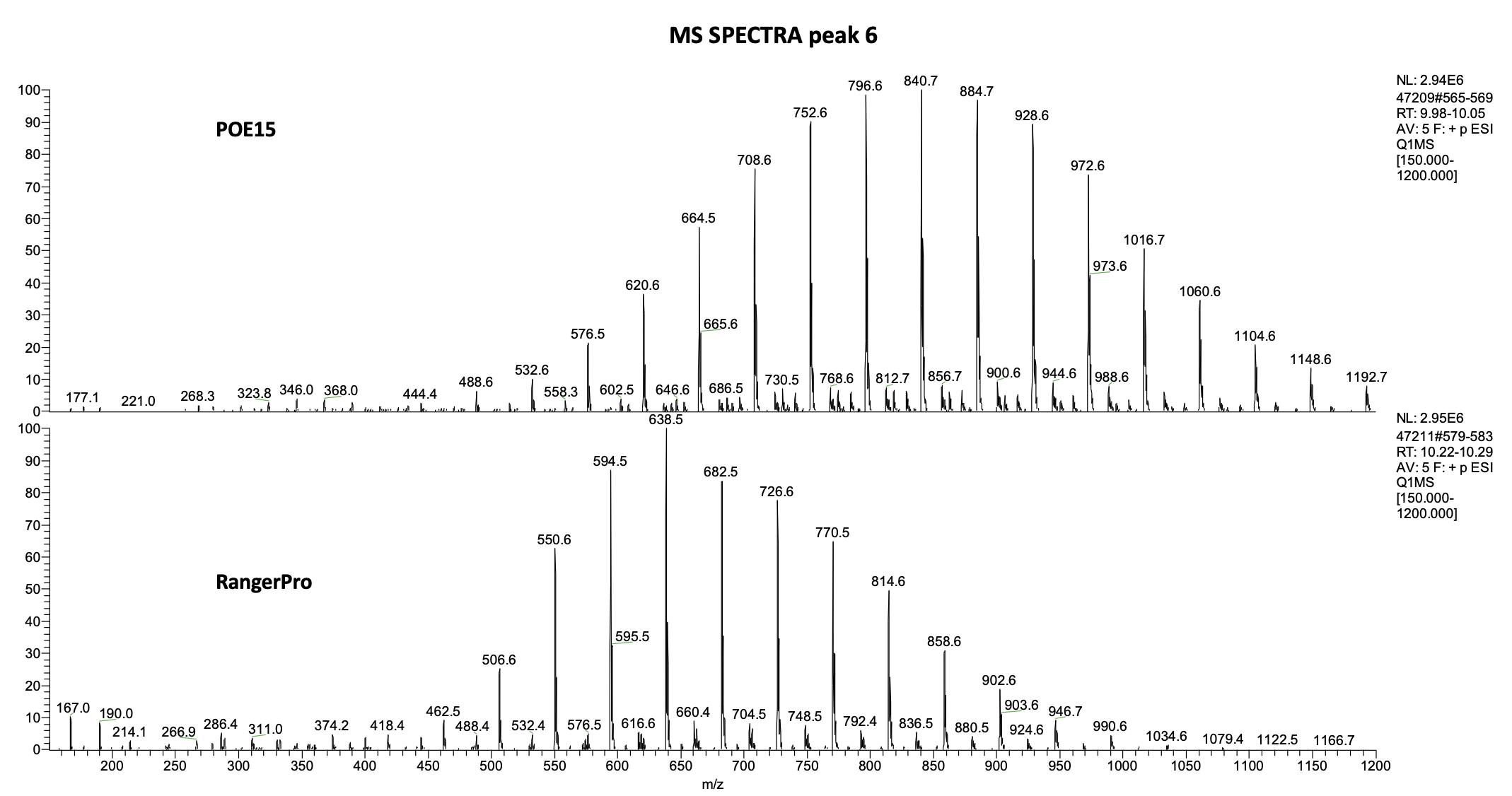

*Representative mass spectra obtained by extracting the TIC of peak 6 from the chromatogram of POE-15 and RangerPro which elute at different retention times (See figure 1).*

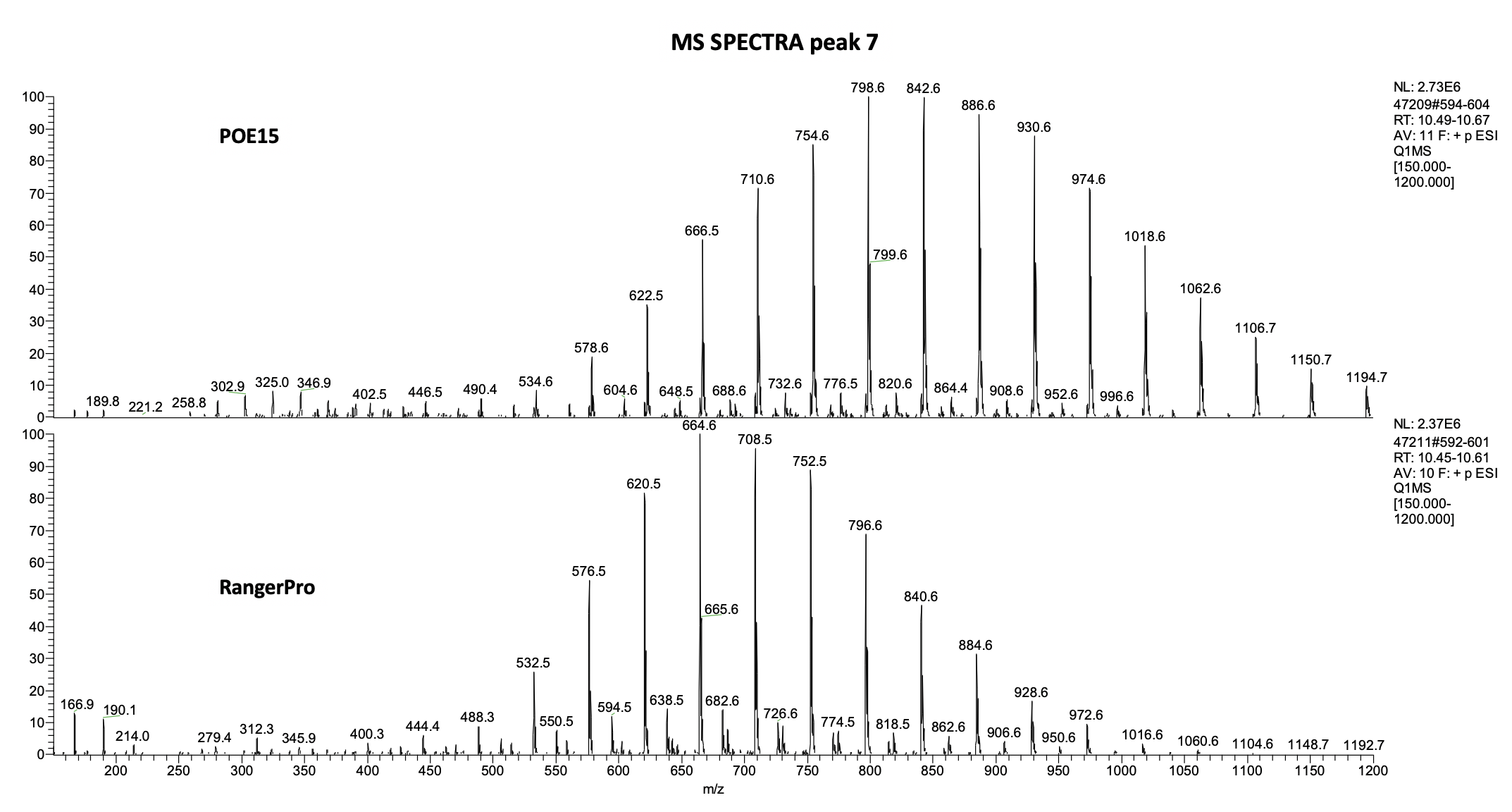

*Representative mass spectra obtained by extracting the TIC of peak 7 from the chromatogram of POE-15 and RangerPro which elute at different retention times (See figure 1).*

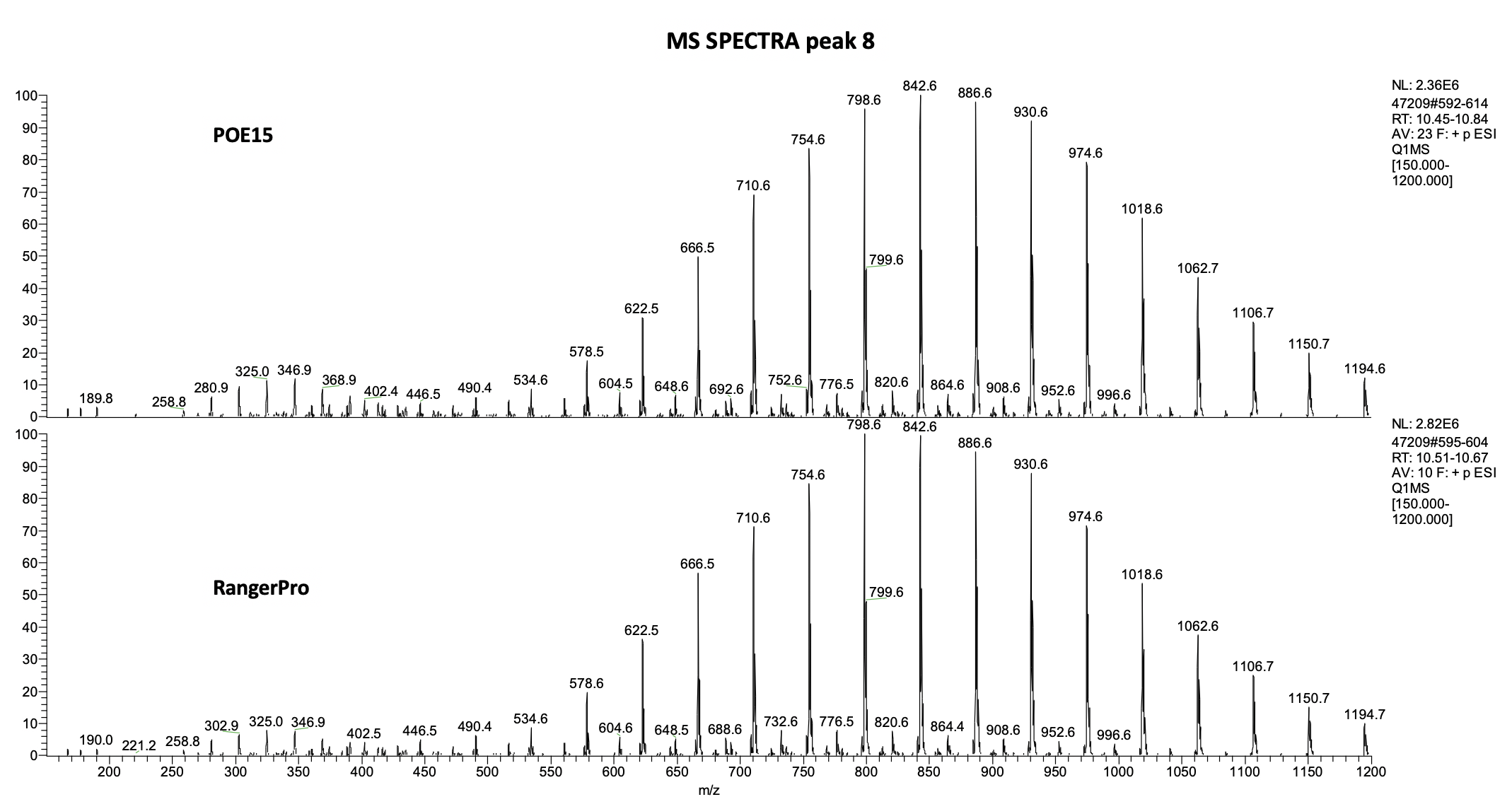

*Representative mass spectra obtained by extracting the TIC of peak 8 from the chromatogram of POE-15 and RangerPro which elute at different retention times (See figure 1).*

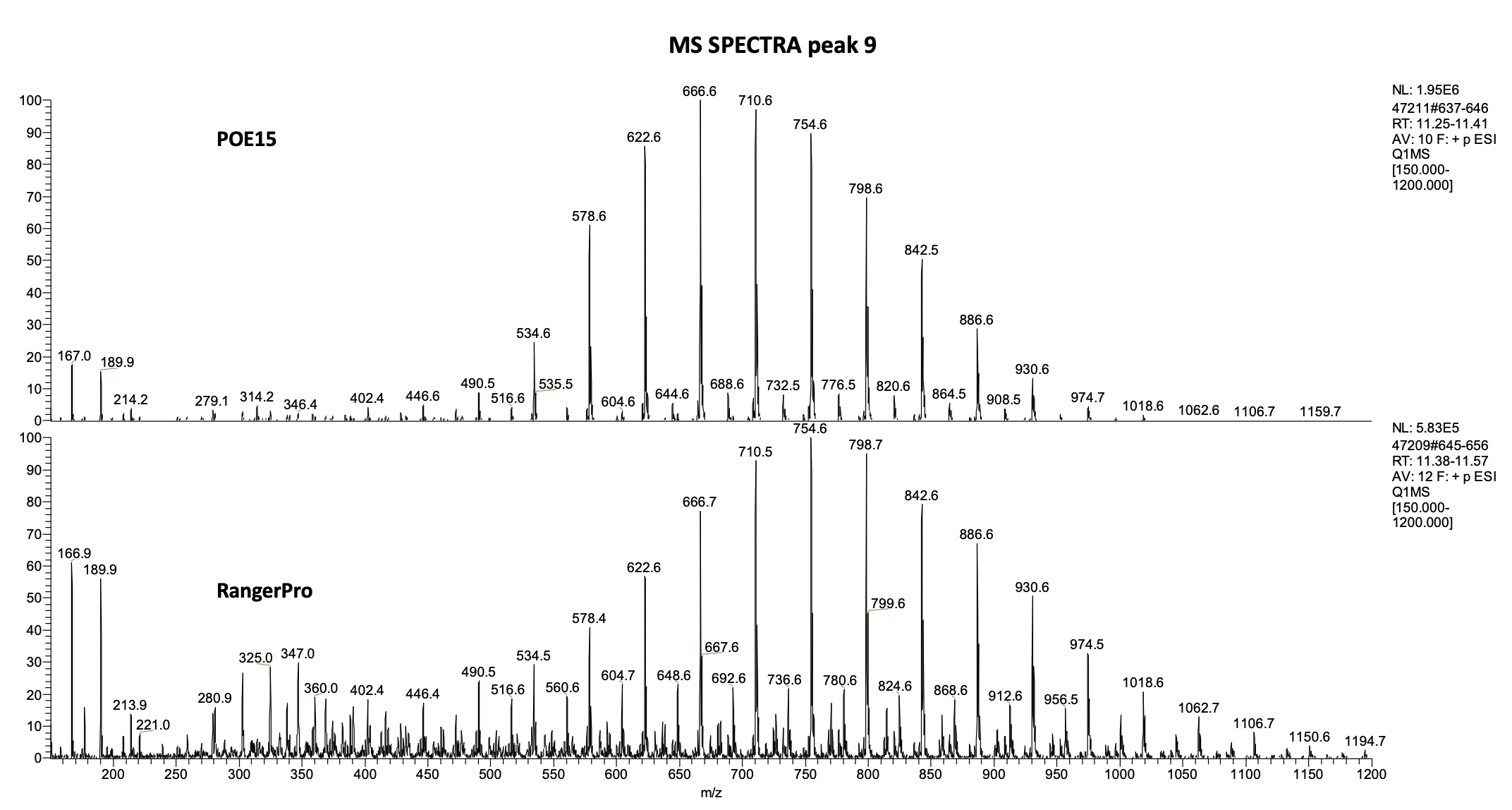

*Representative mass spectra obtained by extracting the TIC of peak 9 from the chromatogram of POE-15 and RangerPro which elute at different retention times (See figure 1).*
